# Multi-Omics Unveils Strain-Specific Neuroactive Metabolite Production Linked to Inflammation Modulation by *Bacteroides* and Their Extracellular Vesicles

**DOI:** 10.1101/2024.08.19.608651

**Authors:** Basit Yousuf, Walid Mottawea, Galal Ali Esmail, Nazila Nazemof, Nour Elhouda Bouhlel, Emmanuel Njoku, Yingxi Li, Xu Zhang, Zoran Minic, Riadh Hammami

## Abstract

*Bacteroides* species are key members of the human gut microbiome and play crucial roles in gut ecology, metabolism, and host-microbe interactions. This study investigated the strain-specific production of neuroactive metabolites by 18 Bacteroidetes (12 *Bacteroides*, 4 *Phocaeicola*, and 2 *Parabacteroides*) using multi-omics approaches. Genomic analysis revealed a significant potential for producing GABA, tryptophan, tyrosine, and histidine metabolism-linked neuroactive compounds. Using untargeted and targeted metabolomics, we identified key neurotransmitter-related or precursor metabolites, including GABA, L-tryptophan, 5-HTP, normelatonin, kynurenic acid, L-tyrosine, and norepinephrine, in a strain- and media-specific manner, with GABA (1-2 mM) being the most abundant. Additionally, extracellular vesicles (EVs) produced by *Bacteroides* harbor multiple neuroactive metabolites, mainly GABA, and related key enzymes. We used CRISPR/Cas12a-based gene engineering to create a knockout mutant lacking the glutamate decarboxylase gene (*gadB*) to demonstrate the specific contribution of *Bacteroides finegoldii*-derived GABA in modulating intestinal homeostasis. Cell-free supernatants from wild-type (WT, GABA+) and Δ*gadB* (GABA-) provided GABA-independent reinforcement of epithelial membrane integrity in LPS-treated Caco-2/HT29-MTX co-cultures. EVs from WT and Δ*gadB* attenuated inflammatory immune response of LPS-treated RAW264.7 macrophages, with reduced pro-inflammatory cytokines (IL-1β and IL-6), downregulation of TNF-α, and upregulation of IL-10 and TGF-β. GABA production by *B. finegoldii* had a limited impact on gut barrier integrity but a significant role in modulating inflammation. This study is the first to demonstrate the presence of a myriad of neuroactive metabolites produced by *Bacteroides* species in a strain- and media-specific manner in supernatant and EVs, with GABA being the most dominant metabolite and influencing immune responses.

**Importance:** *Bacteroides* is a keystone gut symbiont that largely influences gut ecological dynamics and intestinal homeostasis. While previous studies highlighted the contribution of *Bacteroides* to human health, the mechanisms by which these species interact with the gut-brain axis are still emerging. This study highlights the remarkable potential of *Bacteroides* species to produce a wide spectrum of neurotransmitter-related or precursor metabolites, such as γ-aminobutyric acid (GABA), L-tryptophan, 5-hydroxytryptophan (5-HTP), tyramine, normelatonin, L-tyrosine, norepinephrine, and spermine. *Bacteroides* neurometabolic signaling to the host may involve extracellular vesicles (EVs), potentially modulating the gut-brain axis and host immune responses. Notably, *B. finegoldii* exhibit distinct anti-inflammatory characteristics resulting from different molecular patterns, including GABA and EV production. Our findings suggest that *Bacteroides* and their EVs hold great promise as next-generation psychobiotics.

## 1. Introduction

*Bacteroides* are important members of the human gut microbiome and play significant roles in shaping gut ecology, and influencing host physiology and metabolic processes (1). These obligate anaerobes inhabit the lower intestinal tract of mammals in a stable, long-term, mutualistic association with their hosts (2). Their adaptation to gut ecosystems is facilitated by mechanisms, such as cytochrome *bd* oxidase, which enables them to grow at nanomolar O_2_ concentrations, thereby maintaining low oxygen levels and supporting the growth of other strict anaerobes (3). In addition, *Bacteroides* alter the nutritional landscape of the gut by metabolizing complex carbohydrates via polysaccharide utilization loci (PULs) (1). This process provides simple carbohydrates and essential growth factors such as γ-aminobutyric acid (GABA), supporting the expansion of enteric microbes (4).

Extracellular vesicles (EVs) from *Bacteroides* have recently emerged as a communication pathway between gut microbiota and their hosts, participating in glycan utilization and modulating immune responses (5). For instance, *Bacteroides uniformis*-secreted EVs modulate macrophage responses toward a pro-inflammatory phase (6), whereas EVs from *Bacteroides thetaiotaomicron* promote a healthier immune response mediated by dendritic cells. EVs exert their effects by delivering bioactive molecules into host cells, modulating signaling pathways, and regulating intestinal immunity and homeostasis (7, 8). Recent evidence suggests that EVs might harbor neurotransmitters or their precursors (9), potentially acting as mediators of the gut-brain axis, signaling and impacting both immune responses and brain physiology.

The contribution of *Bacteroides* to human health is associated with neuroactivity and brain function as part of the microbiota-gut-brain axis (4). For example, oral administration of *B. fragilis* has been shown to reduce gut permeability, microbiome dysbiosis, and behavioral abnormalities in a mouse model of autism spectrum disorder (ASD), highlighting the potential of microbial interventions to modulate neurological disorders (10). Moreover, *Bacteroides*, a major GABA-producing genus in the gut (4, 11), has been linked to higher levels of serotonin and myoinositol, which are pivotal for signaling between the enteric and central nervous systems (12). The relative abundance of *Bacteroides* has also been negatively correlated with depression-associated brain signatures (4), providing correlative evidence for the role of microbiome-secreted GABA in brain function. Mason et al. (13) also reported depletion of *Bacteroides* in depression and anxiety. Recent findings from the “Lunar Palace 365” experiment demonstrated that *B. uniformis* can reduce chronic stress in crew members by regulating short-chain fatty acids (SCFAs) and amino acid metabolism (14). *B. uniformis* also reduces the elevated intestinal permeability and IL-1β pro-inflammatory response caused by chronic unpredictable mild stress in rats (14). Evidence suggests a potential role for GABA in modulating immune cell activity under various inflammatory conditions (15). For instance, GABAergic signaling is reduced in patients with ulcerative colitis (16). GABA may reduce inflammation through its inhibitory effect on P38 MAPK signaling (17, 18). Moreover, GABA promotes colon health by inducing SCFAs production (19). Restoring GABA levels in the colon can attenuate colitis by inhibiting the production of pro-inflammatory cytokines such as tumor necrosis factor-α (TNF-α), interferon-γ (IFN-γ), and interleukin-12 (IL-12), while increasing the expression of the anti-inflammatory cytokine IL-10 in colon tissue (20). It has been investigated that GABA also regulates pro-inflammatory macrophage responses, thereby limiting IL-1β production (21).

Recently, we isolated 18 Bacteroidetes strains with variable GABA production capacity in EVs and culture supernatants. For instance, a GABA-producing *Bacteroides finegoldii* released EVs with a high GABA content (4 µM) compared to *Phocaeicola massiliensis* (22), suggesting that the neuroactive production capacity of this group of bacteria is strain specific. Our study focused on the neuroactive potential and strain specificity of *Bacteroides* species and aimed to (i) decipher the production of neurotransmitter-related compounds using comparative multi-omics approaches in both culture supernatants and EVs, (ii) assess the cargo capacity of neurometabolites, such as GABA, in secreted EVs, and (iii) investigate the contribution of GABA production by *Bacteroides finegoldii* in modulating intestinal barrier integrity and LPS-mediated inflammatory responses using a newly developed knockout mutant lacking glutamate decarboxylase (*gadB*), a key gene involved in GABA production, created through CRISPR/Cas-based gene editing.

## 2. Materials and Methods

### 2.1. Bacterial isolates and strain identification using 16S rRNA amplicon sequencing

Strains were isolated from a fecal sample collected from a healthy female donor in Ottawa, Canada, with the approval from the University of Ottawa Research Ethics Board (certificate H-02-18-347) as recently described (23). DNA was extracted using a NucleoSpin Microbial DNA Kit (Macherey-Nagel). The 16S rRNA gene was amplified by PCR using the universal bacterial primers 8F and 1391R (23). The purified PCR products were sent for Sanger sequencing at the DNA sequencing facility in the Ottawa Hospital Research Institute. Sequences were quality-checked using BioEdit, analyzed with the RDP classifier at a 99% similarity threshold for species-level identification, and validated using BLAST in NCBI (24).

### 2.2. Whole genome sequencing and bioinformatics analysis

Genomic DNA extraction, library construction, and genome sequencing of 18 Bacteroidetes strains were done as recently described by Yousuf et al. (23). Phylogenomics was conducted using the GToTree software to extract 90 single-copy marker genes from each genome using HMMER (25, 26). Sequences were aligned with MUSCLE, trimmed with TrimAl, filtered to exclude genes with a median length of less than 20%, and a phylogenetic tree was constructed using FastTree (v2.1.11) based on the maximum likelihood method (27) and visualized using iTOL (v.6). Metabolic pathways, including neurotransmitter pathways, were identified using the Pathway Tools software (v. 28) (28) with PGAP-annotated genomes. Genomic, metabolomic, and proteomic figures were visualized by ‘R’ (v4.3.1) (https://www.R-project.org/) in the RStudio 2023.06.0 Build 421 environment using ggplot2 (v3.5.1) (29), corrplot (v0.92) (30), and pheatmap (v1.0.12) (31) packages. A PERMANOVA with 999 permutations was conducted using the Adonis2 function in the vegan package (v2.6.6.1) (32).

### 2.3. Extraction of cell-free supernatant (CFS) and extracellular vesicles (EVs)

Bacterial cells were grown in Fastidious Anaerobic Broth (FAB) supplemented with 10 mM glutamic acid or MacFarlane medium (MFM; and incubated at 37°C for 72 h in an anaerobic chamber. MFM is a complex nutritional medium that closely replicates the nutrients found in the large intestine of healthy adults (33). CFS was obtained by centrifuging at 12,000 × g for 10-30 minutes and filtering the supernatant through a 0.22 µm filter. EVs were isolated from CFS by ultracentrifugation at 100,000 × g for 1 h at 4°C (34). The EV pellet was resuspended in PBS and stored at −80°C until use.

### 2.4. Nanoparticle tracking analysis (NTA) of EVs

EVs isolated from Bacteroidetes strains were characterized using NTA to determine their size distribution and concentration. 1:1000 diluted EVs were analyzed using the ZetaView system (Particle Metrix GmbH, Meerbusch, Germany), measuring vesicles at 11 distinct camera positions. Data were subsequently analyzed using ZetaView software (v8.02.28).

### 2.5. Quantification of SCFAs using gas chromatography

CFSs from selected isolates grown in FAB (+10 mM glutamate) or MFM were analyzed using gas chromatography (Shimadzu-GC-2030; Japan) equipped with a Stabilwax-DA capillary column (60 m x 0.25 mm; Restek) and a flame ionization detector (FID) (34). All analyses were performed in triplicates.

### 2.6. Untargeted and targeted metabolomics by nano-flow LC–MS/MS

To 100 μL of Bacteroidetes CFS and EVs, 900 μL of methanol (−20°C) and 1 µL of 1 mg/mL reserpine (internal standard, Sigma-Aldrich) were added, and the mixture was vortexed for 1 min. The mixture was kept at −20°C for 15 mins, thawed, at room temperature for 3 min, vortexed again, and centrifuged at 12,000 × g for 5 min at 4°C. The supernatant was collected and dried using the SpeedVac system. The dry extract was dissolved in 100 μL 50% acetonitrile, 50% water, and 0.1% formic acid (positive mode). The samples were filtered through a 0.2 μm PTFE filter, transferred into LC vials, and loaded onto a nanoLC coupled to a Q-Exactive Plus mass spectrometer (Thermo Fisher Scientific). Metabolites were separated using a Proxeon EASY nLC II System with a Thermo Scientific Acclaim PepMap RSLC C18 column.

For targeted metabolomics, neuroactive metabolites were identified by matching m/z signals and retention times with commercial pure standards, including GABA, glutamate, tryptophan, 5-HTP, kynurenic acid, normelatonin, l-tyrosine, norepinephrine, dopamine, tyramine, spermine. A 6-point calibration curve was generated, ranging from 1 to 1000 μM for each standard. Peaks with a signal intensity of less than 2500 or less than twice the average blank signal were removed. The raw spectra were processed and deconvoluted using ZMine 2.53. Tandem MS scans were analyzed using SIRIUS by comparing the top 5 candidates from the PubChem organic formula database. Molecular candidates were retrieved, fragmented *in silico*, and matched with experimental spectra. The top candidates for each variable were cross-referenced using the HMDB (https://hmdb.ca) and MiMeDB (https://mimedb.org/). PCA and heat maps were generated using MetaboAnalyst (https://www.metaboanalyst.ca).

### 2.7. Proteomes analysis using nano-flow LC–MS/MS

Protein contents from CFS and EVs were quantified using Bradford assay (#23200, Thermo Fisher Scientific) following manufacturer’s protocol. Samples containing 50 μg of proteins were added in a lysis buffer (25 mM HEPES, pH 8.0, 8M urea, 0.5 % DDM, 1 mM DTT, and 5% glycerol) with a volume ratio of 5:1 (fraction: buffer) and incubated for 15 mins. The mixture was transferred to a 10 kDa filter (MRCPRT010, Millipore Sigma). A filter-aided sample preparation (FASP) method was used for the protein reduction and alkylation with TCEP and iodoacetamide (34). Filter-aided sample preparation (FASP) was used for the reduction and alkylation with TCEP and iodoacetamide. After protein reduction and alkylation, buffer exchange was performed using a digestion buffer (0.6% glycerol and 25 mM HEPES, pH 8.0), followed by proteolytic digestion with trypsin at an enzyme: protein ratio of 1:300 and shaking at 500 rpm at 37°C for 12 h. Digested protein samples were desalted using TopTip C-18 columns and vacuum-dried using the SpeedVac system. The dry peptides were reconstituted in 20 μL of water with 0.1% formic acid and analyzed using an Orbitrap Fusion mass spectrometer coupled with an UltiMate 3000 nanoRSLC (Thermo Fisher Scientific, Mississauga, ON, Canada) (34). The data were processed and searched against the UniProtKB Reference Proteome database using MaxQuant v1.6.2.352 (35). Identified proteins were functionally annotated using eggNOG mapper (http://eggnog-mapper.embl.de/).

### 2.8. Genome editing of Bacteroidetes finegoldii to obstruct GABA biosynthesis

To inhibit GABA biosynthesis in *B. finegoldii*, we deleted the glutamate decarboxylase gene (*gadB*) using a modified version of the method described by Zheng et al., 2022 (36). The aTc-inducible CRISPR/FnCas12a plasmid system (PB025, ∼12 kb) from Addgene (PB025 #183092) was engineered to target *gadB*. Primers were designed using Primer3 (v. 0.4.0)-amplified regions flanking *gadB* (B3/B4 and B5/B6 primer pairs) from *B. finegoldii* genomic DNA. A 25 nt sgRNA targeting *gadB* was designed using CHOPCHOP (https://chopchop.cbu.uib.no/) to ensure that the PAM sequence was 5’-TTN-3’ or TTTN, thus minimizing the off-target effects. Plasmid DNA was prepared using a Plasmid DNA Miniprep Kit (Qiagen, Hilden, Germany). The PB025 plasmid was linearized and *gadB*-directed sgRNA was introduced using the B1/B2 primer pair (**Table S1**) and Phusion High-Fidelity PCR Master Mix (Thermo Scientific™). The residual circular plasmid was digested with DpnI (NEB). The linearized plasmid containing sgRNA (∼10 kb) was purified and fragments flanking *gadB* were ligated using Gibson assembly (NEB). The primer and sgRNA sequences are listed in **Table S1**.

#### Transformation and Conjugation

The Gibson-assembled plasmid (∼11 kb) was transformed into *E. coli* S17-1 λPIR-competent cells, and clones were selected on LB plates containing 100 μg/mL ampicillin. *E. coli* S17-1 was conjugated with *B. finegoldii* in BHIS broth, plated on BHIS agar, and incubated under anaerobic conditions at 37°C for 24 h. The bacterial lawn was resuspended in BHIS medium and serial dilutions were plated on BHIS agar containing gentamicin (200 μg/mL) and erythromycin (25 μg/mL). Colony PCR was performed to confirm the presence of this plasmid in *B. finegoldii*.

#### Construction of ΔgadB Mutant: Anhydrotetracycline-Mediated Induction of CRISPR-Cas

*B. finegoldii* with the PB025 plasmid was grown overnight at 37°C in BHIS with gentamicin and erythromycin under anaerobic conditions. The culture was diluted 1:100 and induced with aTc (100 ng/mL) for 24 h. The serial dilutions were plated on BHIS aTc agar and incubated at 37°C for 48 h. Mutants were confirmed by colony PCR and sequencing. The PB025 plasmid was cured by culturing the mutant in BHIS without antibiotics for 24 h and passaging 10 times, with 100-fold dilutions each time.

### 2.9. Impact of CFS from B. finegoldii wildtype vs mutant (ΔgadB) in an in vitro gut model of epithelial integrity

Caco-2 (passage 10) and HT-29 (passage 16) cells were obtained from ATCC (VA, USA) and co-cultured in HyClone High Glucose Dulbecco’s Modified Eagle’s Medium (DMEM, with L-glutamine) (Cytiva, MA, USA) supplemented with 10% fetal bovine serum and 1% (v/v) non-essential amino acids. The co-culture was seeded at a density of 10^5^ cells in a 9:1 (Caco-2) ratio on Transwell clear inserts with 0.4 µm pore polyester membranes (Corning, NY, USA) and incubated for 28 days at 37°C in a 5% CO₂ atmosphere. The culture medium was refreshed every 2–3 days. This co-culture forming a functional epithelial barrier (37), and was exposed to 10 μg/mL LPS from *Serratia marcescens* (Sigma) to mimic leaky gut conditions. The impact of CFS from *B. finegoldii* wild-type (WT) and Δ*gadB* mutant on monolayer cell integrity was assessed by measuring transepithelial resistance (TEER). Additionally, the Lucifer yellow permeability test (100 μM, Sigma-Aldrich) (38) was performed by adding Lucifer yellow to the apical compartment of the Transwell. The percentage passage of Lucifer Yellow from the apical to the basolateral compartment was determined by measuring the fluorescence at an excitation/emission wavelengths of 485/535 nm.

### 2.10. Impact of CFS and EVs from B. finegoldii wildtype vs mutant (ΔgadB) on macrophage mediated immune response

RAW264.7 macrophages (ATCC TIB-71) were seeded at 10^5^ cells/well in 96-well plates, cultured for 24 h, and subsequently treated with 5 µg/mL LPS (39). CFS and EVs from *B. finegoldii* wildtype and Δ*gadB* mutant strains were added, and the cells were incubated for 24 h. After incubation, cells were harvested for assays to measure the GSH/GSSG ratio, ATP levels, and RNA extraction. Oxidative stress was assessed by measuring the GSH/GSSG ratio using the GSH/GSSG-Glo™ assay (Promega, Madison, USA) (40) and cell viability was determined by measuring ATP levels using the CellTiter-Glo® assay (Promega, Madison, USA) (40) following the manufacturer’s instructions. Luminescence was measured using a Tecan plate reader. Controls included bacterial media alone (negative control), GABA (positive control), and RAW264.7 cells without LPS treatment (background control).

### 2.11. Cytokine gene expression using qRT-PCR

Cells were collected from 96-well plates in triplicate, homogenized with TRIzol reagent (Thermo Fisher Scientific) for RNA extraction, and purified using an miRNA kit (Qiagen). RNA quality was assessed using NanoDrop spectrophotometer and gel electrophoresis. cDNA was synthesized using an iSCRIPT cDNA Synthesis Kit. Quantitative real-time PCR was performed using the SYBR Green Master Mix on a Bio-Rad CFX96 system. Gene expression levels were normalized to those of GAPDH and β-actin. Primer sequences for inflammatory cytokines (TNF-α, IL-1β, and IFN-γ), anti-inflammatory cytokines (IL-10, TGF-β, and IL-6), and housekeeping genes are listed in Supplementary **Table S1**. Data were analyzed using the ΔΔCt method to quantify relative gene expression levels (41).

### 2.12. Statistical analyses

Statistical tests and figure generation were performed using GraphPad Prism 9.0 (GraphPad, San Diego, CA, USA) and R software (v4.3.1). The results are expressed as the mean ± Standard Error. Statistical differences between the study groups were determined using one-way and two-way analyses of variance (ANOVA). Differences were considered statistically significant when the *p*-value was less than 0.05.

## 3. Results

### 3.1. Genome-based metabolic profiling reveals a neuroactive potential in human gut Bacteroidetes strains

Initially, we compared 18 isolates at the genomic level via whole-genome sequencing of each strain. The draft whole genomes of the sequenced strains were clustered into 3 different clusters: *Parabacteroides* spp., *Phocaeicola* spp., and *Bacteroides* spp. (**Fig. 1**). The total number of functional pathways detected across the strains ranged from a minimum of 210 in the two isolates of *Bacteroides stercoris* (UO.H1039 and UO.H1035) to a maximum of 252 in *Phocaeicola vulgatus* UO.H1015 (**Fig. 1**). The number of metabolites detected of the main pathway categories across the 18 strains is presented in **Table S2**. Next, we aimed to identify which genome encodes a predicted neuroactive-related function, including tyrosine, tryptophan, glutamate, phenylalanine, tyramine, norepinephrine, dopamine, and melatonin metabolism. In all 18 isolates, we predicted genes related to L-glutamate, GABA, L-tryptophan, L-tyrosine, and L-phenylalanine metabolism (**Fig. 1**). Moreover, all genomes, except *B. finegoldii* UO.H1052, included genes involved in tryptamine and serotonin metabolism. While all isolates related to *Parabacteroides* and *Phocaeicola* harbored genes involved in dopamine metabolism, only four isolates, *B. Zhangwenhongii* UO.H1054, *P. dorei* UO.H1033, *P. vulgatus* UO.H1015, and *P. vulgatus* UO.H1016, had norepinephrine-related metabolic pathways (**Fig. 1**).

**Figure 1.**
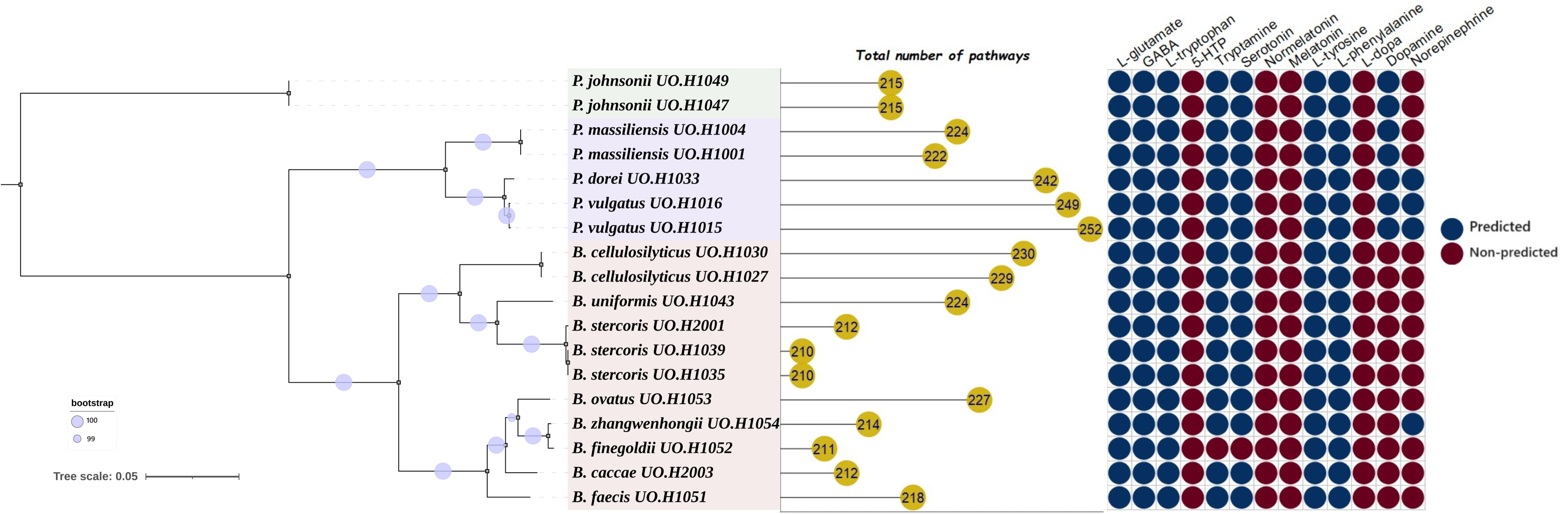
Neurotransmitter pathways distribution across the 18 neuroactive gut bacterial strains based on WGS data. A phylogenomic tree was constructed with GToTree software and visualized with iTOL. Blue circles indicate the bootstrap percentage of 1000 replicates. Pathway numbers were identified relying on the *de novo* assembled genomes. The last panel is the neurotransmitters and their precursors, as presence/absence, were extracted from the pathways data generated using pathway tools software.

### 3.2. Bacteroidetes strains have distinct neurometabolism

#### 3.2.1. Bacteroidetes strains produce distinct neuroactive metabolome profiles

To explore the neuroactive metabolite production potential of *Bacteroidetes*, we conducted untargeted metabolomics of CFS from 18 strains after 72h of fermentation. In each strain, we identified 150-175 extracellular metabolites, including amino acids, lipids, organic acids, adenosines, glutathione, coenzymes, sugars, and nucleotides (**Fig. S1; Table S3**). The identified metabolites included a diverse array of neuroactive molecules such as GABA, Spermine, 5-HTP, and dopamine (**Fig. 2A**). All tested isolates produced high levels of GABA, with glutamate decarboxylase—a key enzyme in GABA synthesis—being one of the most abundant exoproteins identified across all 18 strains (**Fig. S2; Table S4**). Tryptophan and its derivatives, 5-HTP, normelatonin, and kynurenic acid, were detected at lower levels (**Fig. 2A**). For example, 5-HTP is produced by *P. johnsonii* UO.H1047, UO.H1049, *P. massiliensis* UO.H1001, *P. vulgatus* UO.H1016, *B. stercoris* UO.H1035, and *B. faecis* UO.H1051. Dopamine was detected in four isolates (*P. dorei* UO.H1033, *B. stercoris* UO.H1035 and UO.H1039, and *B. zhangwenhongii* UO.H1054).

**Figure 2.**
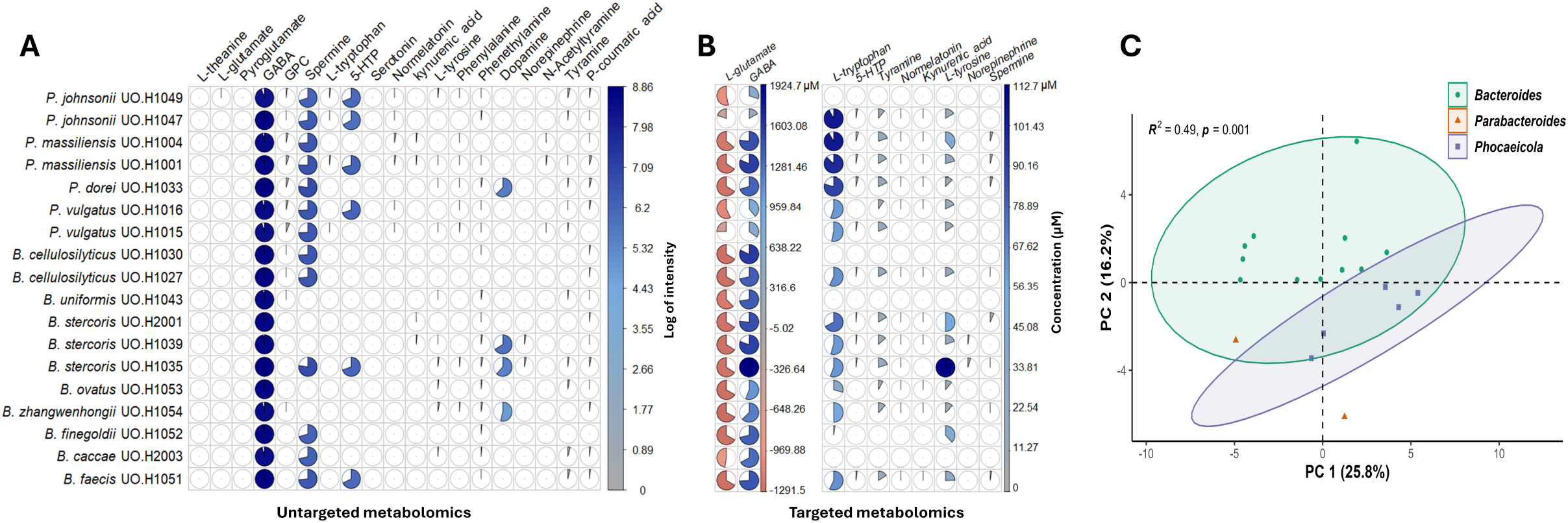
Detection of neurotransmitter-related neuroactive metabolites. (A) The untargeted metabolomics approach shows the abundance of numerous potential neuroactive metabolites released by Bacteroidetes strains. (B) Illustrates the differences between the 18 strains in producing different neuroactive compounds and their precursors as quantified by targeted metabolomics analysis. (C) Bacteroidetes strains showed separate clusters based on genus indicating that producing neuroactive metabolites is a strain-specific trait as shown by PCA plot depending on the targeted metabolomics data. *R*² is the coefficient of determination and *p* refers to the *p*-value.

To validate the findings of untargeted metabolomics, we conducted targeted quantification of various neuroactive metabolites, including GABA, L-glutamate, L-tryptophan, 5-HTP, normelatonin, kynurenic acid, L-tyrosine, dopamine, norepinephrine, spermine, and tyramine (validation presented in **Fig. S3**). Using a 6-point calibration curve (1–1000 μM, R² > 0.99), we detected high concentrations of GABA (0.3 - 2 mM) in all the isolates, and to a lesser extent, the accurate masses of L-tryptophan (15–120 μM), L-tyrosine (12–110 μM), and tyramine (10-20 μM) (**Fig. 2B**). Normelatonin, spermine, norepinephrine, and 5-HTP were quantified in certain isolates at concentrations ranging from 0.1 to 10 μM, whereas kynurenic acid was detected at the lowest concentrations (0.025–0.125 μM). Dopamine was not detected in any of the samples possibly because of its low concentration or false positive identification with Sirius software. Principal component analysis (PCA) of targeted neurometabolite concentrations revealed that the genus was a key differentiating factor in the metabolic profiles (**Fig. 2C**).

#### 3.2.3. Bacteroides EVs harbor a distinct array of neuroactive metabolites

GABA production was also compared between *P. massiliensis* UO.H1001, *B. faecis* UO.H1051, *B. finegoldii* UO.H1052, and *B. ovatus* UO.H2003 strains in both SUP and EVs using ELISA. The results showed that *B. finegoldii* UO.H1052 released the highest GABA concentrations with approximately ∼6.5 mM in SUP and ∼4.0 µM EVs. Strains UO.H1051 and UO.H2003 exhibited similar GABA levels, whereas the lowest concentrations were observed in *P. massiliensis* UO.H1001, both in SUP and EVs (**Fig. 3A**).

**Figure 3.**
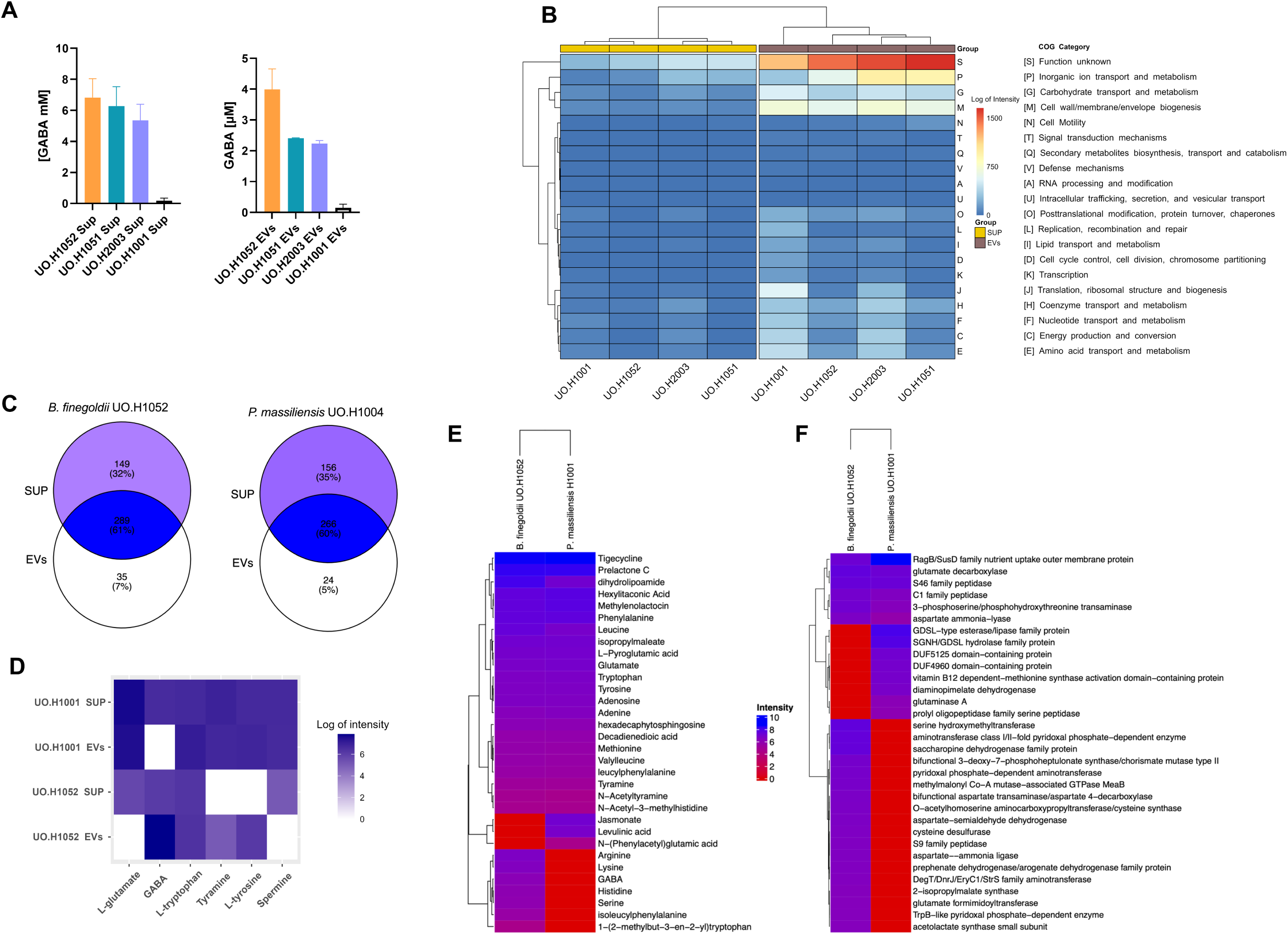
Comparative metabolomic and proteomic analyses of SUP and EVs between selected Bacteroidetes strains grown on FAB medium. (A) A Venn diagram of untargeted metabolites detected in SUP and EVs of 4 strains. (B) An interactive cluster heatmap comparison between SUP and EVs shows the COG categories of 4 strains. (C) Bar plots illustrate GABA production comparison between the 4 strains in SUP and EVs, *B. finegoldii* UO.H1052 and *P. massiliensis* UO.H1001 grown on (FAB) exhibited the highest and lowest GABA levels respectively. (D) A heatmap shows the abundance of some neuroactive metabolites produced by strains UO.H1052 and UO.H1001. (E) A heatmap explained the production of neuroactive metabolites based on untargeted metabolomics analysis while (F) is a heatmap that identified the protein profile for UO.H1052 and UO.H1001.

Proteomic analysis revealed distinct protein profiles across the 4 strains, varying between SUP and EVs (Fig 3B). EVs from strain UO.H1001 contained the highest number of proteins (>800), followed by UO.H2003 (>700), whereas UO.H1052 and UO.H1051 EVs contained over 500 identified proteins (**Table S5**). Functional analysis clustered the identified proteins into 20 clusters of orthologous groups (COGs). The most common protein functions in EVs are inorganic ion transport and metabolism (P), cell wall/membrane/envelope biogenesis (M), and carbohydrate transport and metabolism (G). However, a large proportion of proteins in all 4 EV samples were classified as having unknown functions (S) (**Fig. 3B**).

A comparative analysis between the highest GABA producer (*B. finegoldii* UO.H1052) and the lowest producer (*P. massiliensis* UO.H1001) revealed distinct metabolic profiles of the SUP and EVs of these two strains, as determined by untargeted metabolomics (**Fig. 3C; Table S6**). Further metabolomic and proteomic analyses of strains UO.H1001 and UO.H1052 showed diverse source-specific (SUP or EVs) and strain-specific production of neuroactive compounds (**Fig. 3D**). Multiple compounds and their precursors were identified within EVs, such as GABA, L-glutamate, phenylalanine, L-tryptophan, L-tyrosine, and tyramine (**Fig. 3D**). Additionally, the EVs of *B. finegoldii* UO.H1052 were uniquely characterized by the presence of arginine, histidine, lysine, and serine amino acids (**Fig. 3E**). Proteomic analysis further revealed that glutamate decarboxylase was highly abundant in the EVs of both bacterial strains, whereas glutaminase A was only detected in the EVs of UO.H1052 (**Fig. 3F**).

#### 3.2.2. SCFA production is strain and media specific

We quantified SCFA concentrations in the CFS of 18 isolates grown on FAB and MFM media. In FAB, propionic acid was produced at 4 mM concentrations in most strains, except *P. massiliensis* UO.H1001 and UO.H1004, where acetic acid (∼3-4 mM) was predominant. *B. caccae*, *B. faecis*, *B. finegoldii*, *B. ovatus*, and *B. zhangwenhongii* produced the lowest SCFA concentrations compared to other strains (<2 mM). In MFM, SCFAs were produced in significantly higher amounts than in FAB medium. For example, acetic acid was produced at concentrations >30 mM in most strains, except for *B. caccae*, *B. faecis*, *B. finegoldii*, *B. stercoris* UO.H1039, *B. uniformis* UO.H1043, *P. vulgatus* UO.H1015, *P. vulgatus* UO.H1016, *P. johnsonii* UO.H1049, and *P. massiliensis* UO.H1001, which showed lower acetic acid concentrations (10-15 mM). Propionic and butyric acids were higher in all isolates than in FAB, with the highest levels (∼10 mM) found in *P. massiliensis* UO.H1004, *P. dorei*, *B. ovatus*, and *B. zhangwenhongii* (**Fig. 4A-B**). PCA indicated that SCFA production was medium- and genus-specific (**Fig. 4C-D**).

**Figure 4.**
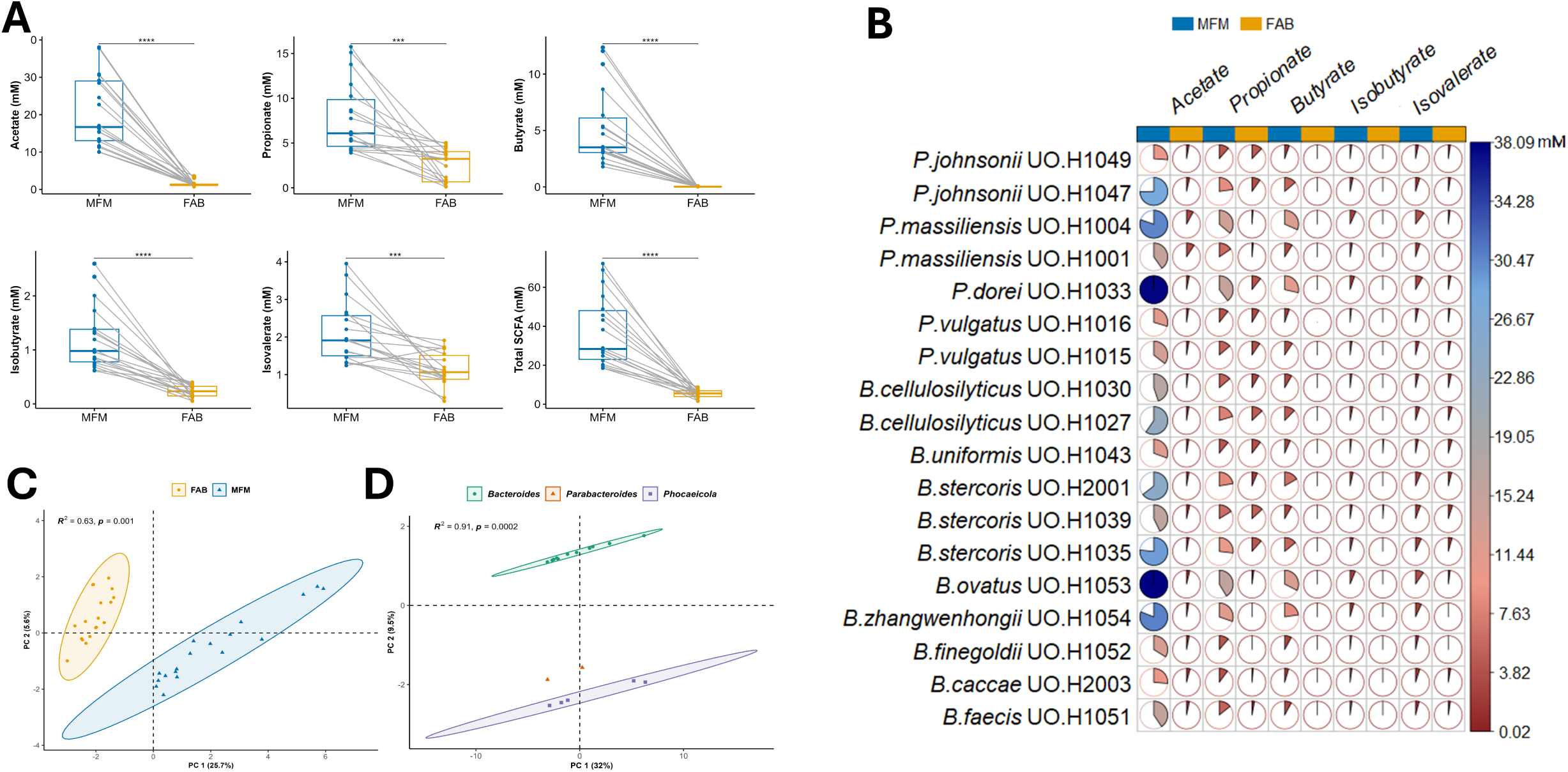
Comparative quantification of SCFAs production by Bacteroidetes strains in MacFarlane (MFM) and FAB media. Paired plots revealed the significant impact of fermentation media on the production of different SCFAs between MFM and FAB media. (B) Pie plots show the variation in concentrations of SCFAs released by each strain relying on the fermentation medium. (C&D) PCA plots showing the clustering of Bacteroidetes strains in groups based on the growing medium (C) or the taxonomy level (D). \*\*\**p <* .001, \*\*\*\**p <* .0001. *R*² is the coefficient of determination and *p* refers to the *p*-value.

### 3.3. Genome editing of B. finegoldii UO.H1052 to Create a ΔgadB knockout mutant

Genome analysis revealed that these 18 Bacteroidetes strains possessed a highly conserved glutamate decarboxylase (GAD) system, including glutamate decarboxylase (*gadB*), glutaminase (*glnA*), glutamate/GABA antiporter (*gadC*), and potassium channels. We selected *gadB* from the *B. finegoldii* genome for knockout to assess its impact on GABA production. The *gadB* gene, consisting of 1437 bp, was successfully deleted using an inducible CRISPR/FnCas12a approach (**Fig. 5A, B**), and this deletion was confirmed by polymerase chain reaction (PCR), agarose gel electrophoresis, and sequencing (**Fig. 5C**). The Δ*gadB* deletion mutant exhibited a reduced growth rate compared with its wild-type (WT) counterpart when cultured in minimal medium with xylose as the sole carbon source (**Fig. 5D**). Using a targeted metabolomics approach based on LC-MS/MS, GABA production in the WT supernatant was measured to be ∼800 µM, whereas the Δ*gadB* deletion mutant exhibited no GABA production in the micromolar range (**Fig. 5D**). Disruption of the GABA production phenotype in the Δ*gadB* mutant further validated this knockout. Additionally, glutamate levels were ∼50 µM in the WT and 1500 µM in the Δ*gadB* mutant (**Fig. 5D**), indicating that GABA was generated via the decarboxylation of glutamate.

**Figure 5.**
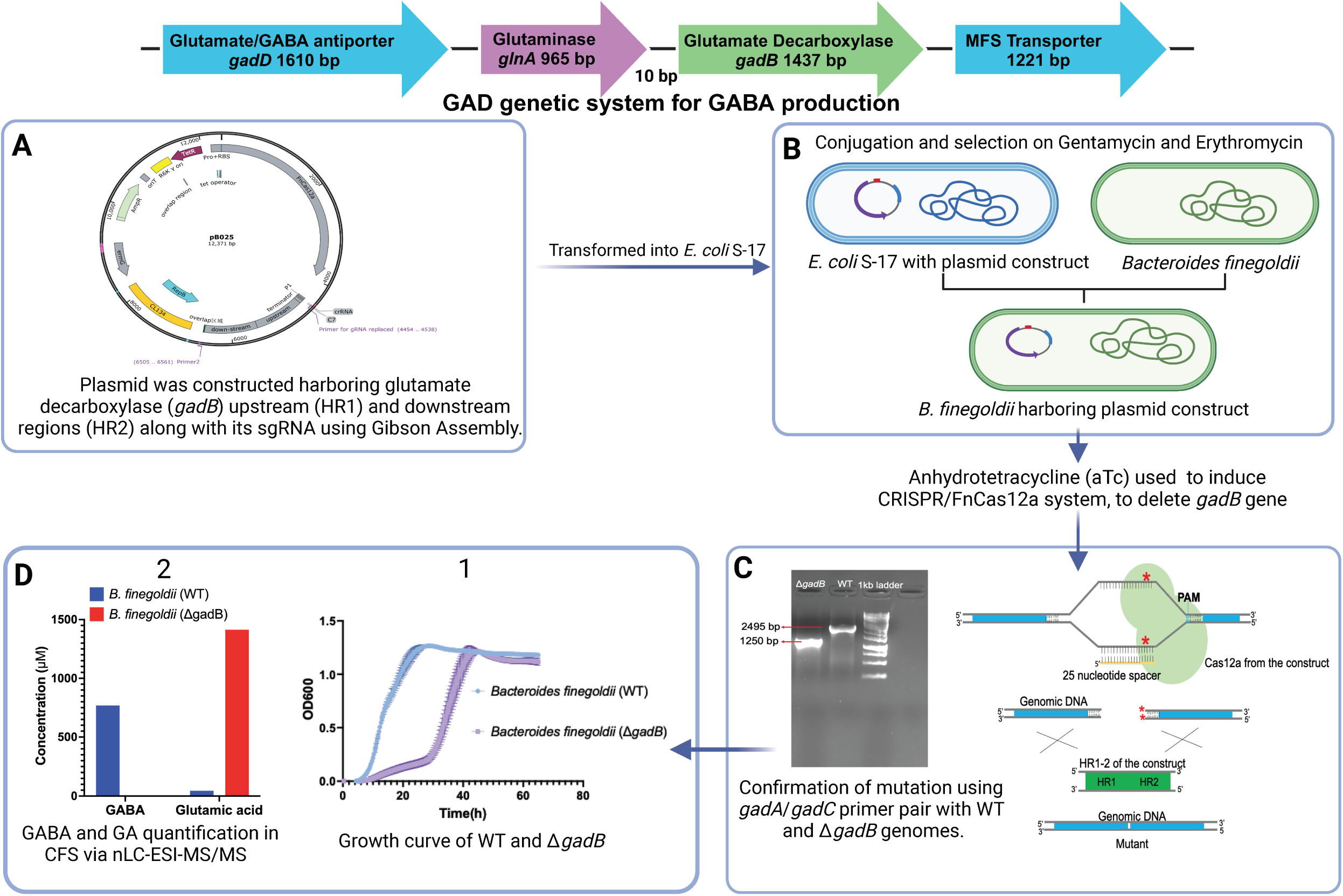
Obstruction of GABA biosynthesis by knocking out the glutamate decarboxylase gene (Δ*gadB*) using the CRISPR/FnCas12a system. (A) The PB025 plasmid was constructed by introducing sgRNA and upstream and downstream flanking regions of the *gadB* gene. (B) The constructed plasmid was transformed into *E. coli* S17-1, and then conjugated into *B. finegoldii*. (C) The expression of the Cas protein was induced with anhydrotetracycline (aTC) to activate the deletion of the *gadB* gene and confirmed by PCR and sequencing. (D-1) shows the differences in the growth curve between *B. finegoldii* WT and mutant strain (Δ*gadB*) in a minimal medium with xylose. (D-2) A bar plot illustrates the variations in GABA and glutamic acid production levels in WT and Δ*gadB* strains at stationary phase CFS relying on targeted metabolomics analysis.

### 3.4. GABA production by Bacteroides finegoldii: Limited impact on gut barrier integrity, Significant role in inflammation

To evaluate the effect of GABA production by *B. finegoldii* UO.H1052 on gut barrier integrity, we developed a Δ*gadB* mutant (GABA-) lacking GABA production (**Fig. 5C**). We compared the effects of the WT (GABA+) and Δ*gadB* (GABA-) strains on the integrity of Caco2/HT29-MTX cell monolayers treated with LPS (5 μg/mL). Transmembrane electrical impedance (TEER) was measured pre-exposure, post-LPS exposure at 24 h, and post-treatment at 48 h. TEER levels at 24 h significantly decreased from 280±5 to 260±5 Ω·cm² in all LPS-treated groups, except for the non-LPS control. At 48 h, both WT and Δ*gadB* CFS significantly increased the TEER to 330±7 Ω·cm² in the presence of LPS (*p* < 0.00251). The TEER in the DMEM+LPS group decreased to 250±3 Ω·cm² (*p <* 0.01), whereas the TEER in the DMEM-only group remained stable at 280±5 Ω·cm². The addition of GABA did not enhance TEER, which remained at 260 Ω·cm². No significant differences were observed between the WT and Δ*gadB* CFS treatment groups (**Fig. 6A**). The Lucifer yellow assay confirmed the modulation of paracellular permeability. LPS treatment significantly increased permeability (*p <* 0.0001) compared to the negative control. Both WT and Δ*gadB* CFS treatments markedly reduced Lucifer yellow crossing into the basolateral compartment (*p <* 0.0001), indicating superior membrane integrity compared with the DMEM+LPS, GABA+LPS, and NC+LPS groups (**Fig. 6B**). EVs extracted from *B. finegoldii* and Δ*gadB* analyzed via NTA, revealed that 95% of these vesicles fell within the size range of approximately 137–188 nm, with a concentration of 7±3 × 10^9^ particles/mL (**Fig. S4**). To assess the effects of GABA production on host inflammatory responses, RAW macrophages were stimulated with LPS (5 µg/mL) followed by treatment with CFS or EVs from WT, Δ*gadB*, or pure GABA (500 µM) (**Fig. S5**). After 24 h, the samples were analyzed for oxidative stress, ATP activity, and cytokine gene expression. EVs from WT *B. finegoldii* and pure GABA induced the highest GSH/GSSG ratios (14.5±1 and 16±1, respectively). Although Δ*gadB* EVs increased the GSH/GSSG ratio compared with that of the control, this effect was lower than that of WT EVs (*p <* 0.0002). CFSs from both WT and Δ*gadB* strains did not positively affect cellular redox homeostasis (**Fig. 6C**). The ATP viability assay showed that GABA and WT EVs resulted in the highest ATP levels (*p <* 0.01 and 0.0008, respectively) (**Fig. 6D**). Δ*gadB* EVs did not alter cellular ATP metabolism compared with the control (*p <* 0.05). We also quantified pro-inflammatory (TNF-α, IL-1β, IL-6, and IFNγ) and anti-inflammatory cytokine (TGF-β1 and IL-10) gene expression in RAW macrophages, LPS-exposed RAW macrophages, and LPS-exposed cells treated with EVs from WT, Δ*gadB* and GABA. RAW macrophages treated with LPS compared to RAW macrophages only showed significantly higher pro-inflammatory gene expression (IL-6, ∼60-fold), IL-1β (∼7.5-fold), TNF-α (∼3.5-fold), IFNγ 3.5-fold (**Fig. S5**). GABA or WT EVs downregulated TNF-α (−2.4±0.05 and −2.0±0.2) and upregulated IL-10 (1.30±0.3 and 2.5±0.4) and TGF-β1 (2.0±0.5 and 2.1±0.1). Δ*gadB* EVs significantly increased TNF-α (∼3-fold) and IL-1β (∼4.5-fold) while downregulating IL-6 (∼1.2-fold). WT EVs and GABA treatments also increased IL-1β (∼2-fold), but to a lesser extent than Δ*gadB* EVs, and upregulated IL-6 (∼1.5-fold). IFNγ expression was slightly upregulated (∼1.3-fold) by WT EVs, whereas the other treatments downregulated it by ∼2-fold (**Fig. 6E**).

**Figure 6.**
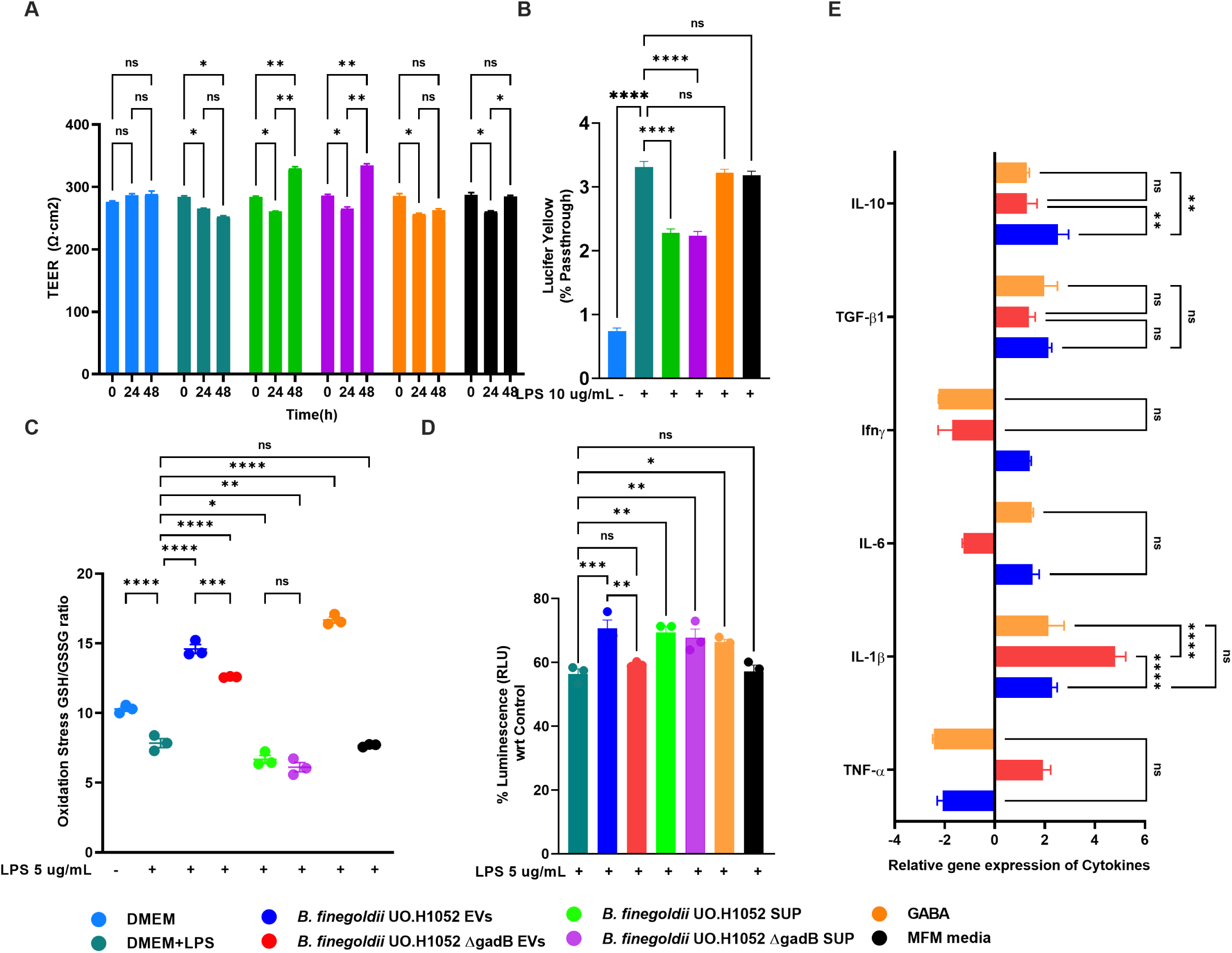
*B. finegoldii* CFS attenuates LPS-induced gut membrane disruption in Caco-2/HT29 co-cultures, while GABA-enriched EVs modulate inflammatory responses in RAW 264.7 macrophages. (A) Exposure of Caco-2/HT29 co-cultures to LPS resulted in a significant decrease in TEER after 24 h. The subsequent treatment with bacterial supernatants of WT and Δ*gadB B*. *finegoldii* strains showed a significant increase in transepithelial resistance. (B) Lucifer Yellow assay showed reduced florescence in wells treated with WT and Δ*gadB*, indicating attenuated leaky gut conditions compared to control groups. (C) LPS-induced oxidative stress in RAW264.7 macrophages was significantly reduced by WT and Δ*gadB* EVs and GABA based on GSH/GSSG ratio measurement. Contrarily, the CFS increased oxidative stress, with Δ*gadB* showing higher oxidative stress than WT and GABA. (D) Δ*gadB* EVs exhibited the lowest cell viability compared to WT EVs and CFS from WT, Δ*gadB*, and GABA, as measured by ATP levels. (E) Gene expression by qRT-PCR analysis with LPS-treated RAW264.7 macrophages showed that WT EVs and GABA downregulated TNF-α and upregulated IL-10 and TGF-β, with IL-10 significantly less upregulated in the ΔgadB mutant. EVs with GABA and chemically synthesized GABA similarly affected IL-1β and IL-6 expression. \**p <* .05, \*\**p <* .01, \*\*\**p <* .001, \*\*\*\**p <* .0001.

## 4. Discussion

*Bacteroides* is a major genus within the human gut microbiome that contributes to shaping gut ecological dynamics (42, 43) and maintaining intestinal homeostasis (1, 44). This study focused on the neuroactive potential and specificity of *Bacteroides* species and examined the spectrum of neurotransmitter-related compounds using a multi-omics approach. We also sought to assess the neuroactive cargo capacity of EVs and elucidate the specific contribution of *Bacteroides*-derived GABA in modulating the gut barrier integrity and LPS-mediated immune responses.

Our multi-omics analyses revealed that 18 Bacteroidetes strains (*Bacteroides* [11 isolates], *Parabacteroides* [2 isolates], and *Phocaeicola* [5 isolates]) produced a distinct array of neuroactive metabolites, including GABA, tryptophan derivatives, and catecholamines. The results support the previous phylogenetic assessments of Bacteroidetes genomes, showing very little difference between *Bacteroides* and *Parabacteroides* species, with a shared core of 1085 protein families (42). Our findings also align with those of Karlson et al. (2011), showing that the genomes of our isolates are enriched in carbohydrate metabolism, amino acid metabolism, and cofactor synthesis (42). However, the metabolic profiles of these closely related species suggest that they may contribute differently to human health. For instance, *B. fragilis*, *B. uniformis*, *B. caccae*, and *B. ovatus* colonization have been reported to have different effects on tryptophan metabolites, including serotonin, along the gut-brain axis (45). All strains encoded for glutamate/GABA, tryptophan, L-tyrosine, and L-phenylalanine metabolic pathways, with norepinephrine and dopamine pathways predicted only in *Parabacteroides* and *Phocaeicola* isolates. *Parabacteroides* spp. have abundant β-glucuronidase activity, which elevates catecholamine levels in the gut (46, 47). Our metabolomic results revealed strain-specific neurometabolite production, with GABA being the most dominant, and to a lesser extent, L-tryptophan, 5-HTP, tyramine, normelatonin, L-tyrosine, norepinephrine, and spermine. This study is the first to describe 5-HTP production in *Bacteroides*, an intermediate in the serotonin biosynthesis pathway, highlighting the unique metabolic capacity of these strains. *Parabacteroides* species enrichment in a mouse model of depression and its end products, tryptophan metabolites, have been correlated with neuroplasticity in a brain region linked to depression (48). Notably, a selective tryptophan metabolite induces antidepressant-like behaviors in mice (48). Normelatonin (N-acetyl serotonin), another tryptophan metabolite that was first reported in this study, is an antioxidant and neuroprotective molecule that suppresses the NF-κB pathway (49). Our findings also show the presence within EVs of multiple neurotransmitter-related compounds and related metabolic pathways, mainly GABA and its related enzymes. EVs protect these molecules from lytic enzymes in the gut environment and facilitate their horizontal transfer across both short (e.g., intestine) and distant (e.g., brain) locations (53–55). In addition, the lipid nature of EVs facilitates their uptake and processing by host cells, enabling them to cross biological barriers, such as the intestinal and blood-brain barriers (22). Taken together, this data demonstrates that *Bacteroides* produce strain- and media-specific neurometabolites with important significance in the gut-brain axis.

Furthermore, *B. finegoldii* metabolome induced a GABA-independent improvement of intestinal barrier integrity, as no significant difference was observed between the WT and *ΔgadB* CFS treatments. Previous administration of a *Bacteroides* strain to mice increased gut barrier integrity and reduced inflammation through induction of Treg cells, IL-22, and SCFA production (53). The anti-inflammatory effect of GABA was evident in our results through the reduction of TNFα expression in a GABA-dependent manner and its synergistic effect on increasing the anti-inflammatory cytokine IL-10 expression in RAW cells. The anti-inflammatory effect of GABA relies on the suppression of T cell responses and reduction of pro-inflammatory cytokines (54, 55). GABA stimulates GABA_A_ receptors on immune cells, leading to the suppression of T1 responses, reduction of NF-κB activation, decrease in pro-inflammatory cytokines, and stimulation of IL-10 signaling (54, 55). In addition, *B. fragilis* attenuates DSS-induced colitis in mice through the TLR2/IL-10 signaling pathway (56). LPS induces inflammation through TLR4, while TLR2 recognizes different molecular patterns, including lipoproteins and glycolipids, and in some reports LPS (57), which could explain the distinct upregulation of pro-inflammatory cytokines observed in our results. In this study, *B. finegoldii* EVs induced GABA-dependent expression of IL-1β. In agreement with this, EVs of the probiotic *L. reuteri* DSM 17938 induced the expression of IL-6 and IL-1β in peripheral blood mononuclear cells (58). This was attributed to the abundance of 5′-nucleotidase (5′NT) in EVs, which converts AMP into the signal molecule adenosine (58). In agreement with this, we detected 5′-NT in the proteome of EVs released by *B. finegoldii. Bacteroides* can also modulate inflammation, cytokine levels, and gut ecology through propionate production (44). Overall, this data indicated that *B. finegoldii* possesses distinct anti-inflammatory characteristics resulting from different molecular patterns, including GABA and EV production.

In summary, this study highlights the potential of *Bacteroides* species as key gut microbes for producing a broad spectrum of neurotransmitter-related compounds such as GABA, tryptophan derivatives, and catecholamines. Our findings suggest an important role for *Bacteroides* neurometabolism in modulating the gut-brain axis and host immune responses. *Bacteroides* signaling to the host involves EVs, which transport several neuroactive metabolites. Although further research is needed to decipher the microbiome-host dialogue and how these gut microbes affect host neurobiological functions, *Bacteroides* and their EVs hold great promise as next-generation psychobiotics. However, their safety and efficacy as probiotics requires further investigation.

## Data availability

The 16S rRNA gene sequence and genome data of the bacterial isolates are available from the NCBI database (<u>OP690542-OP690599</u> and <u>PRJNA898401</u>, respectively). The mass spectrometry proteomics data were deposited in the ProteomeXchange Consortium via the PRIDE partner repository with the dataset identifiers <u>PXD054243</u> and <u>PXD045162</u> (Usernames; Passwords: hTODkNLHFcbD and 1CFQzgzm, respectively). Metabolomic data were deposited in the EMBL-EBI MetaboLights database using the identifier <u>MTBLS10526</u>.

## Consent for publication

All authors consent to the publication.

## Author contributions

B.Y., R.H., and W.M. conceived of and designed the study. B.Y. and N.E.B. isolated the gut bacteria and performed molecular identification. G.A.E., W.M., B.Y., and R.H. analyzed the genomic data and performed the bioinformatics analysis. B.Y., Y.L., E.N., and Z.M. prepared and processed the samples for the metabolomic and proteomic studies. B.Y., and Z.M. analyzed the metabolomic data. B.Y. conducted the CRISPR/Cas study. B.Y. and N.N. conducted cell culture experiments. B.Y., X.Z., and W.M. analyzed the proteomics data. B.Y., W.M., G.A.E., and R.H. wrote or revised the manuscript. All authors reviewed and approved the final manuscript.

## Acknowledgements

We thank Dr. Ayman ElSayed for technical assistance. We are grateful to Dr. Linggang Zheng and Dr. Lei Dai from the Shenzhen Institute of Synthetic Biology, Shenzhen Institutes of Advanced Technology, Chinese Academy of Sciences, Shenzhen 518055, China, for making the CRISPR/FnCas12a plasmid accessible through Addgene and providing essential tips for genome editing of *B. finegoldii*. We would like to thank undergraduate students Ana PR Muñoz, Charlie Levine, and Joseph SW Mariam, as well as master’s student Mariem Chiba, for their assistance with various experiments. Mass spectrometry was performed at the John L. Holmes Mass Spectrometry Facility at the University of Ottawa.

## Supplementary Figures

**Figure S1.** Comparative untargeted metabolomics among 18 Bacteroidetes strains. (A) Principal Component Analysis (PCA) of all metabolites identified via untargeted metabolomics in positive mode, analyzed using MetaboAnalyst, shows distinct separation among the 18 strains. (B) Heatmap of Log-transformed peak intensities for highly abundant metabolites by untargeted metabolomics analysis.

**Figure S2.** Glutamate decarboxylase is among the highly abundant exoproteins identified in all 18 strains. Heatmap showing LFQ intensity of highly abundant exoproteins across 18 Bacteroidetes strains. In contrast to glutamate decarboxylase, glutaminase A was not identified in all strains. Histidine and aspartate decarboxylases were detected exclusively in specific strains.

**Figure S3.** Validation of the identification and quantification of neuroactive metabolites and/or their precursors from CFS of 18 *Bacteroidetes* strains based on extracted ion chromatography peaks with a mass tolerance of <5 ppm. The peaks were selected based on retention time, and m/z values of the pure standard compounds.

**Figure S4.** Nanoparticle tracking analysis (NTA) of the EVs extracted from *B. finegoldii* UO.H1052 and its derived Δ*gadB* mutant. (A) Representative image of EVs detected from a NTA video. (B) Size distribution of EVs revealed 95% EVs are in the average size range of 160 nm.

**Figure S5.** Gene expression of anti-inflammatory and pro-inflammatory cytokines in RAW cells relative to LPS-treated RAW cells. The data were normalized to the expression levels of the housekeeping genes GAPDH and β-Actin.

## Supplementary Tables

**Table S1.** RT-qPCR primer sequences of genes were analyzed in this study for expression analysis.

**Table S2.** Comparison of main pathway categories across 18 strains with the number of compounds detected per pathway.

**Table S3.** Comprehensive metabolite profiling of CFS from 18 Bacteroidetes strains isolated from the human gastrointestinal tract via untargeted metabolomics.

**Table S4.** Exoproteome profiling of CFS from 18 Bacteroidetes strains based on nano-LC-MS.

**Table S5.** Proteomics of EVs extracted from *P. massiliensis* UO.H1001, *B. faecis* UO.H1051, *B. finegoldii* UO.H1052, and *B. ovatus* UO.H2003. Functional annotation was done with eggNOG mapper (http://eggnog-mapper.embl.de/) for all the proteins.

**Table S6.** Untargeted metabolomics of EVs extracted from *P. massiliensis* UO.H1004 and *B. finegoldii* UO.H1052 strains.

